# Computational deconvolution of fifteen leukocyte subtypes from DNA methylation microarrays trained on flow cytometry data in the Health and Retirement Study

**DOI:** 10.1101/2022.11.28.518190

**Authors:** Jonathan A. Heiss, Kelly M. Bakulski, Bharat Thyagarajan, Eileen M. Crimmins, Jessica D. Faul, Jonah D. Fisher, Allan C. Just

**Author notes:** < >, < >, < >, < >, < >, < >, < >.

## Abstract

Adjusting for cell composition is critical in epigenome-wide association studies of whole blood samples. Using DNA methylation of whole blood samples (as opposed to purified cell types) and complete blood counts/flow cytometry data from 2530 participants in the Health and Retirement Study, we trained and tested a computational model that extends the number of estimated leukocyte subtypes to fifteen compared to established models with six or seven cell types. Our model, which can be applied to both Illumina 450k and EPIC microarrays, explained a larger proportion of the observed variance in whole blood DNA methylation levels than popular reference-based cell deconvolution approaches, and vastly reduced the number of false-positive findings in a reanalysis of an epigenome-wide association study of chronological age.

## Introduction

Epigenome-wide association studies (EWAS) search for biomarkers of past or current exposures, lifestyle habits, or diseases. The majority of EWAS measure DNA methylation (DNAm) in whole, unfractionated blood samples [1]. Processing blood for cell type fractionation requires special treatment during sample collection and the additional logistics and costs often render such workflows infeasible for large-scale epidemiological studies. Yet cell composition is a major source of variance of whole blood DNAm and can influence many of the associations reported in recent years, such as age, smoking, BMI, and birth weight [2]. Consequently, a lack of directly measured cell proportions is often discussed as a major limitation. This conundrum is resolved by computational methods that estimate cell proportions from the whole blood DNA methylome that are used in turn to adjust for these unobserved confounders. The most commonly applied methods provide estimates of six or seven major cell lineages and are based on reference DNA methylomes from purified cell types [3,4]. Criticism persists, however, whether such adjustment is sufficient, or whether varying proportions of subtypes continue to confound EWAS results. For rare or closely related subtypes, purifying sufficient quantities to generate reference DNA methylomes is technically challenging, which is a limitation of cell deconvolution models that rely on purified end-members of each cell type. The aim of this project was to leverage a uniquely large dataset with both whole blood DNAm and complete blood counts/flow cytometry measures to implement a computational deconvolution algorithm that does not require fractionated methylome references and extends the repertoire of cell types for which predictions are possible.

## Methods

### Study sample

The United States Health and Retirement Study (HRS) is a nationally representative panel cohort of Americans over the age of 50 [5]. Beginning in 1992, the HRS was conducted every two years through interviews, surveys, and biosampling. In the 2016 Venous Blood Study of the HRS [6], biosamples were collected in participants’ homes by trained phlebotomists. Included in the venous blood panel was a 10mL EDTA tube processed as whole blood for DNAm experiments and complete blood counts (CBC), and an 8 mL cryopreservation tube (CPT) for later flow cytometry. The CPT was shipped to the University of Minnesota Advanced Research and Diagnostic Laboratory at room temperature while the EDTA tube was shipped at 4°C. Samples were received within 24 hours of collection (92%) or within 48 hours of collection (99%). The CPTs were centrifuged to obtain peripheral blood mononuclear cells, which were cryopreserved using previously published protocols and stored in liquid nitrogen freezers until further use. HRS was approved by the Institutional Review Board at the University of Michigan, Ann Arbor. All HRS study participants provided signed informed consent prior to study participation.

### DNA methylation data cleaning and preprocessing

DNA was isolated from packed cells frozen at −70°C. The DNA extraction and purification method used a modified salt precipitation method for protein removal using commercial Flexigene^®^ reagents (Qiagen Instrument Service, Germantown, MD) and a FlexStar+ Instrument (Autogen, Inc., Holliston, MA). DNA was quantitated using the NanoDrop Spectrophotometer (NanoDrop Technologies, Wilmington, DE). DNA was normalized and assigned to 96-well plates by stratified randomization on key demographic variables (age, cohort, sex, education, race/ethnicity). Bisulfite treatment and cleaning were performed using the Zymo EZ-96 DNA methylation kit, according to the manufacturer’s protocols. DNAm of whole blood samples was assessed on the Infinium MethylationEPIC BeadChip (Illumina, San Diego, CA) according to the manufacturer’s protocols. Samples were excluded if (i) an assay failed according to at least one of 17 control probe metrics; (ii) more than 100,000 of the queried loci were classified as undetected by stringent criteria that we have previously validated [7]; (iii) the sex of the sample donor, as inferred from the signal intensities of probes querying allosomal loci, did not match the records, or if samples did not cluster with the majority of male or female samples; (iv) the β-value distribution of probes querying SNPs indicated the possibility of sample contamination [8] (see Figure 1). Loci that were undetected in more than one-third of all samples were removed. Furthermore, loci were restricted to those common to the MethylationEPIC and HumanMethylation450 BeadChips. Restricting features to this subset allows users to apply the final cell composition model to datasets from either platform. Fluorescence intensities were corrected for dye bias using the within-sample RELIC method [9] but no between-sample normalization was performed. Methylation levels were expressed as β-values, which approximate the linear relation between cell type-specific and whole blood averages.

**Figure 1:**
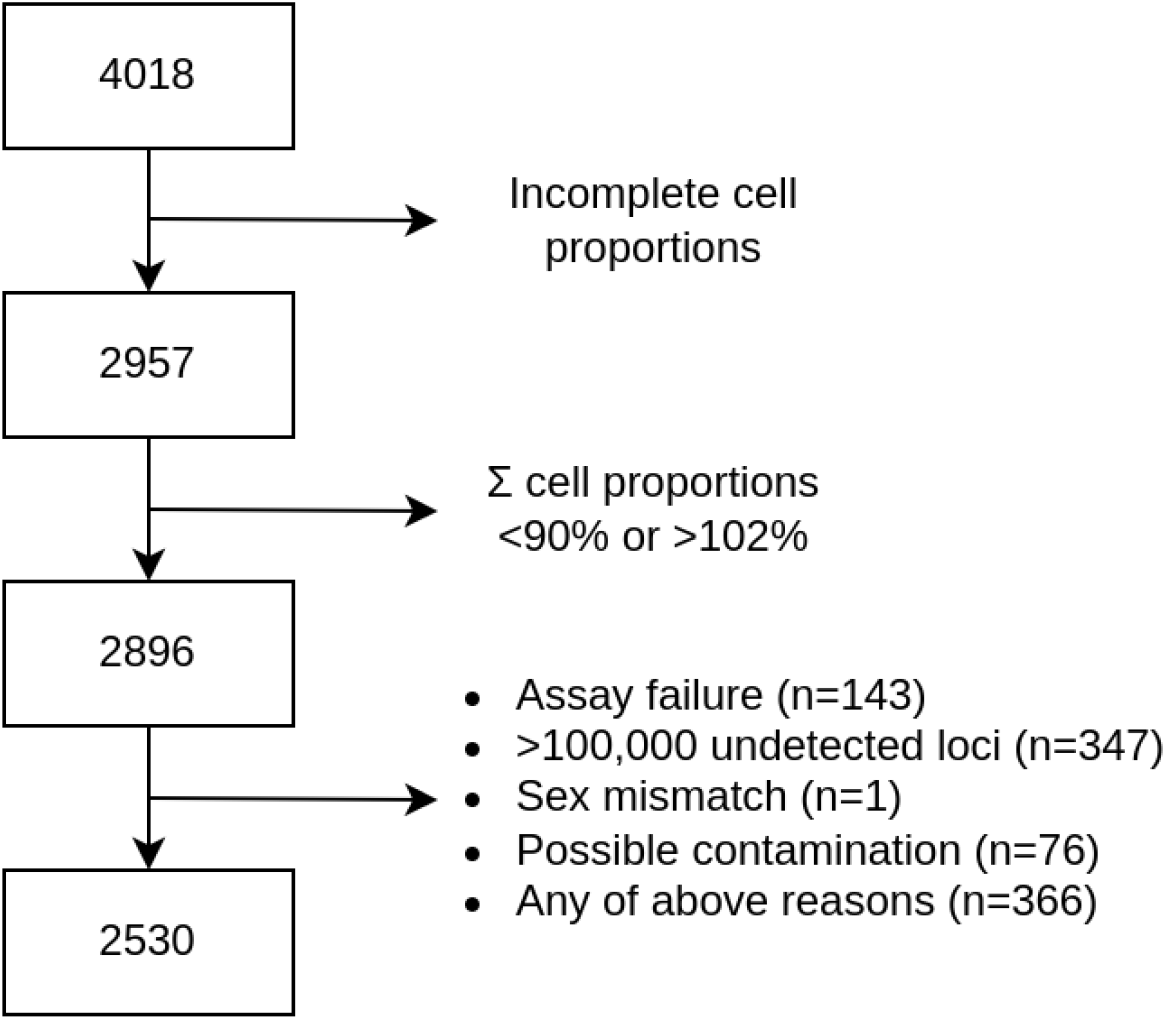
Number of samples passing each filter step. Of the initial 4018 DNA methylation samples, 2530 are available for analysis.

### CBC and flow cytometry preprocessing and cleaning

The granulocyte (neutrophils, eosinophils, basophils) count and monocyte counts were obtained from a complete blood count that was measured in EDTA whole blood using a Sysmex XE-2100 instrument (Sysmex America, Inc., Lincolnshire, IL) within 72 hours after collection.

CPT samples were processed using a standardized protocol as described previously [10,11]. Samples were stained and sorted in two panels. The first panel measured subsets of T cells and B cells and the second panel measured subsets of monocytes, dendritic cells, and natural killer cells [6]. One vial of cryopreserved mononuclear cells containing ~4 million cells was thawed and cells were incubated at 37°C in Roswell Park Memorial Institute media for 1 hour. The cells were centrifuged at 1200rpm for 10 minutes at room temperature. The cells were resuspended in 1× phosphate-buffered saline and stained as outlined previously [12]. The cells were kept on ice until analysis. All flow cytometry measurements were performed on an LSRII flow cytometer or a Fortessa X20 instrument (BD Biosciences, San Diego, CA). This method has been previously shown to estimate cell proportions similar to those obtained from fresh blood [12]. The immunophenotyping data were analyzed using OpenCyto and FlowAnnotator [10].

While the DNA for the microarrays was extracted from whole blood samples, only white blood cells contain nuclei and contribute therefore to the blood DNA methylome (the amount of circulating cell-free DNA is negligible in this context). Hence, only CBC and flow cytometry measurements of cell types belonging to this fraction were included. From CBC we included four cell types: (i) neutrophils, (ii) eosinophils, (iii) basophils, (iv) monocytes. From flow cytometry we included an additional eleven cell types: (v) dendritic cells, (vi) B cells, (vii) natural killer lymphocytes, (viii) naive cytotoxic T cells, (ix) central memory cytotoxic T cells, (x) effector cytotoxic T cells, (xi) effector memory cytotoxic T cells, (xii) naive helper T cells, (xiii) central memory helper T cells, (xiv) effector helper T cells, and (xv) effector memory helper T cells. The selected fifteen cell types are mutually exclusive, i.e., they are not sub/super-types of each other. For example, while we include neutrophils, eosinophils, and basophils – the constituents of granulocytes – the category “granulocytes” itself is not part of the model. At the same time, the list of 15 cell types is exhaustive, meaning that every nucleated cell type present in unfractionated blood is represented. While further subtypes were available in the HRS dataset for some of the select cell types (e.g., counts of myeloid and plasmacytoid dendritic cells) we decided not to use them if doing so would result in proportions close to zero most of the time. More extensive documentation about this HRS dataset (called the 2016 Venous Blood Study) can be found in [6].

Cell type-specific counts (per liter) were divided by the total white blood cell count to convert them to proportions. Samples for which a total could not be calculated because at least one of the 15 cell type proportions was missing, were dropped. As were samples for which the total did not fall between 90-102% (see Figure 1). The lower bound serves to exclude samples with substantial loss during flow cytometry, whereas a violation of the upper bound may indicate errors during data entry or similar.

### Training/testing split

After data cleaning, complete data (complete blood count, flow cytometry, and DNAm) were available for 2530 samples (see Figure 1), of which 1618 were used for model training and 912 were used for testing (the test and training samples were independent). To mitigate the impact of batch effects on our assessment of model performance, batches (i.e., the 96-well plates) were assigned exclusively to either the training or test set, thereby preventing the model from overfitting to batch effects and inflating (internal) model performance. We describe the distributions of participant demographics (sex, age, race, and ethnicity) and total white blood cell count (see Tables 1 and 2).

**Table 1:** Health and Retirement Study participant characteristics in the testing and training datasets.

|  | Training set (n=1,618) | Test set (n=912) |
| --- | --- | --- |
| Sex (male/female) | 644/974 | 397/515 |
| Age in years (median/min/max) | 68/50/100 | 67/51/95 |
| Race (White/Black/other/unknown) | 1211/266/135/6 | 674/161/74/3 |
| Ethnicity (Hispanic/non-Hispanic/unknown) | 247/1370/1 | 127/785/0 |
| White blood cell count in 1e9/L (median/min/max) | 6.4/2.3/15.5 | 6.35/2.7/209 |

**Table 2:** Comparison of estimated with measured cell proportions. All numbers are derived from the test set (n=912). Second column: average measured proportion by flow cytometry; third column: Pearson correlation coefficient between DNA methylation estimated and flow cytometry measured proportions; fourth column: root mean squared error between estimated and measured proportions in percentage points (pp). CBC: Complete blood counts.

|  | Cell type | Measured average (%) | Estimated average (%) | Correlation between measured/estimated proportions | RMSE (pp) |
| --- | --- | --- | --- | --- | --- |
| C<br>B<br>C | Granulocytes |  |  |  |  |
|  | Neutrophils | 57.7 | 60.6 | 0.922 | 4.8 |
|  | Eosinophils | 3.4 | 3.4 | 0.882 | 1.2 |
|  | Basophils | 0.9 | 1.0 | 0.174 | 1.2 |
|  | Monocytes | 8.5 | 8.7 | 0.754 | 1.9 |
| F<br>L<br>O<br>W<br>C<br>Y<br>T<br>O<br>M<br>E<br>T<br>R<br>Y | Cytotoxic T cells |  |  |  |  |
|  | Naive cytotoxic T cells | 1.1 | 1.2 | 0.666 | 0.9 |
|  | Central memory cytotoxic T cells | 0.4 | 0.5 | 0.425 | 0.6 |
|  | Effector cytotoxic T cells | 2.5 | 2.5 | 0.718 | 1.8 |
|  | Effector memory cytotoxic T cells | 1.1 | 1.1 | 0.520 | 1.0 |
|  | Helper T cells |  |  |  |  |
|  | Naive helper T cells | 6.6 | 6.4 | 0.881 | 2.1 |
|  | Central memory helper T cells | 5.3 | 5.2 | 0.750 | 2.2 |
|  | Effector helper T cells | 0.4 | 0.6 | 0.535 | 0.8 |
|  | Effector memory helper T cells | 2.1 | 1.8 | 0.341 | 2.1 |
|  | Natural killer cells | 3.4 | 3.2 | 0.753 | 1.8 |
|  | B cells | 2.3 | 2.4 | 0.721 | 2.5 |
|  | Dendritic cells | 0.8 | 0.8 | 0.176 | 1.0 |

### Feature selection in the training dataset

For this step, the most common cell type, neutrophils, was left out in order to reduce collinearity between input variables. Then, partial Spearman correlation coefficients ρ_it_ were computed, i.e., the correlations between β-values of each locus *i* and the proportions of cell type *t* adjusted for all other cell types (except neutrophils). For each cell type, 50 loci with the highest absolute ρ_it_ were picked. The list of all loci from all cell types, after eliminating any duplicates, constitutes the features of the leukocyte composition model. Loci that were used to estimate cell composition were excluded prior to the evaluation of model performance (again, ensuring independence of training and testing).

### Estimating model parameters in the training dataset and application in the test dataset

β-values of each feature *i* were regressed on the same set of variables as before, i.e., proportions of the fourteen cell types (still without neutrophils). The *lmrob* function implementing robust linear regression from the R package *robustbase* was used [13]. Intercepts α_i_ and regression coefficients β_i_ were recorded to construct a matrix as follows.

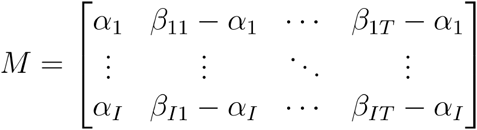

The intercepts, the first column of the matrix, correspond to the neutrophil-specific parameters, i.e., the left-out cell type. Using *M* from the training dataset and the whole blood β-values of the features in the independent test dataset, we estimated cell proportions in the test dataset via quadratic programming (to ensure non-negative proportions) as described by Houseman [4]. Estimates were not constrained to sum up to one.

### Evaluation of model performance in the test dataset

To quantify the share of variance that can be attributed to leukocyte composition alone, but not batch effects or other factors, the following benchmark was designed. Linear regression models were fit for each locus that is autosomal, common to EPIC and its 450K predecessor chip, passed quality control and that was not part of the features utilized to estimate cell proportions. These criteria applied to 437,307 loci. Regression models were fit for the test dataset with four different sets of covariates: the first set **A** included sex, age, batch (96-well plate), and the first five principal components of the matrix of control probe metrics, a similar approach to the one implemented for functional normalization [14]; set **B** extended set **A** by adding the CBC/flow cytometry measured cell proportions, whereas for set **C** the DNAm-derived cell proportions were added instead; finally, set **D** included the covariates from **A** and DNAm-derived proportions of six cell types (granulocytes, monocytes, CD4 T cells, CD8 T cells, B lymphocytes, natural killer lymphocytes) generated by using the same reference dataset [3] as implemented in *minfi*. Coefficients of determination (R^2^) were computed for all linear regression models and the gain in R^2^ from **A** to **B** and **A** to **C** (and from **A** to **D**) was computed and averaged across all loci. The covariates in sets **B** and **C** are of the same number and types, allowing us to compare the R^2^ values and determine which set explains more of the variance associated with leukocyte composition in the test dataset.

### Application to an external dataset

We evaluated our model on a public dataset to demonstrate generalizability. GSE87571, a large (729 samples) dataset from the Gene Expression Omnibus repository featuring a broad age range (14 to 94 years) and using the Infinium HumanMethylation450 BeadChip (including .idat files) was chosen. Our primary covariate of interest was participants’ chronological age, which is well documented for associations with DNAm. The gain in R^2^ for each locus was determined as described above: set **A** included the covariates sex, age, and the first five principal components of the matrix of control probe metrics (information on which 96-well plates samples were allocated was not available); set **C** extended set **A** by adding the estimated cell proportions from our model, whereas for set **D** estimates were derived from the Reinius reference dataset [3]. Furthermore, for each fitted linear regression, the strength of the association with chronological age was recorded and significant loci were counted to get an idea of the magnitude of potentially false-positive findings.

## Results

The HRS training and test datasets had similar demographic characteristics to each other (Table 1). Samples were 60/56% female, 74/74% White, 16/18% Black, and had a median age of 68/67 years, respectively (Table 1). In the test set, the most abundant cell type was neutrophils (57.7%) and the least abundant were central memory cytotoxic T cells and effector helper T cells (both 0.4%) (Table 2).

Pearson correlation coefficients between flow cytometry measured and DNA methylated estimated cell proportions in the HRS test set are listed in Table 2. They range from 0.922 for neutrophils to 0.174 for basophils. In the same test dataset, we compared the variance explained by model **A** without any cell composition adjustment to a series of models with alternative cell adjustment approaches. The average gain in variance explained was 9.5 percentage points for covariate set **B**, which used flow cytometry to measure cell counts. The average gain in variance explained was 14.1pp for set **C**, which used our new DNAm-derived cell estimates. This means that our estimated cell proportions explained on average 4.5pp more than models containing the flow cytometry measured cell proportions. Figure 2 shows the proportion of variance of whole blood DNAm levels explained by measured and estimated cell proportions, respectively, for the top 10,000 loci when ranked by the former metric. Despite this selection of probes that favors measured cell proportions, the estimated cell proportions explained more variance in 9,101 instances. The average gain for set (**D** Reinius reference set) was 10.7pp.

**Figure 2:**
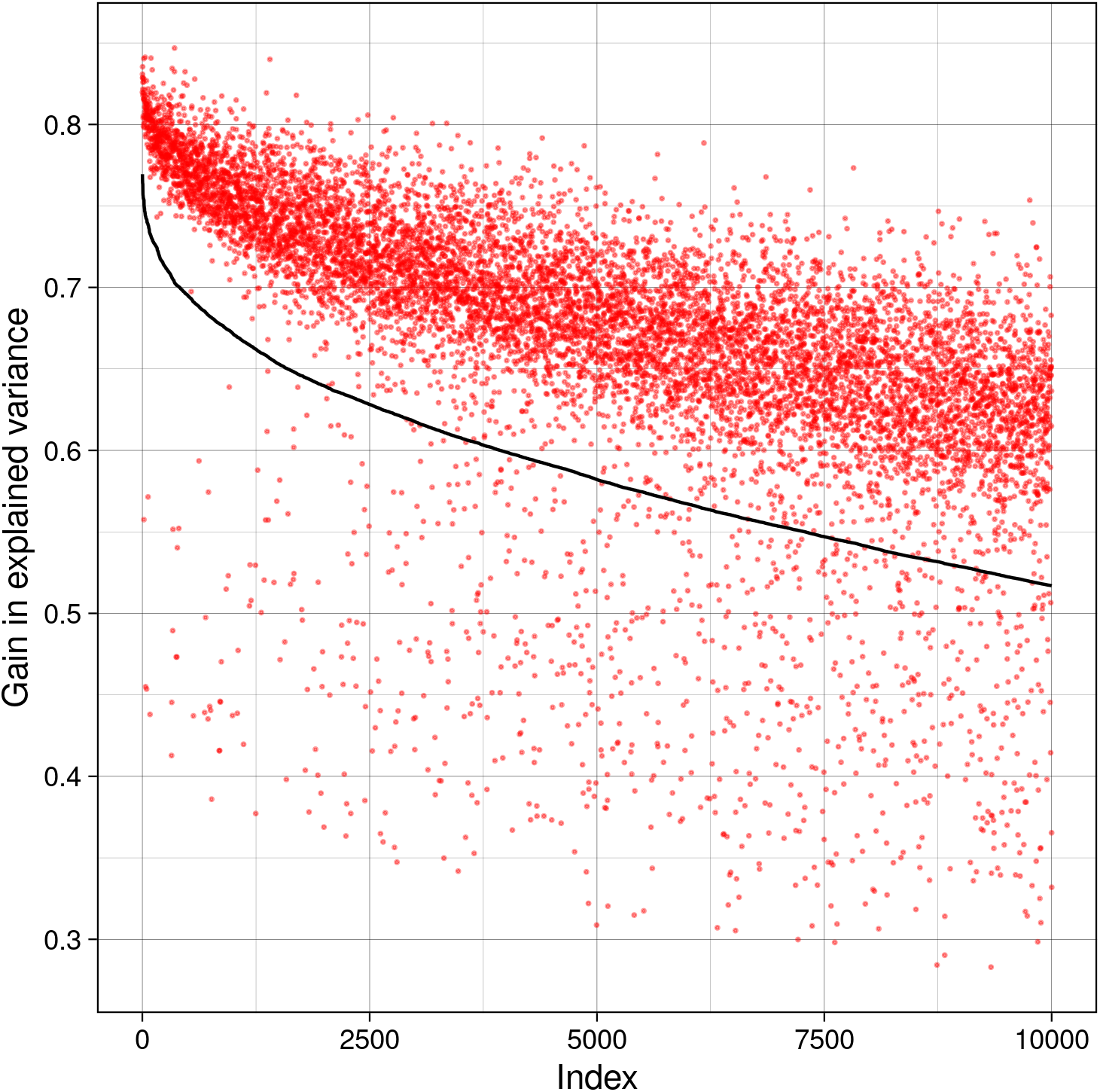
Proportion of variance of whole blood DNAm levels explained by flow cytometry measured (black) and DNA methylation estimated (red) cell proportions, respectively, for the top 10,000 probes when ranked by the former metric.

In the external dataset, compared to models without any cell composition adjustment, models using covariate set **C** with our new DNAm-derived cell proportion estimates had an average gain in variance explained of 18.7pp. For covariate set **C**, we observed 3,029 out of 484,900 loci tested were associated with chronological age at a Bonferroni-corrected significance of 5%. The corresponding numbers for set (**D** adjustment using the Reinius reference panel) were 16.2pp and 120,711 significant loci. Only 87 out of the 3,029 loci were not included among the set of 120,711. Without adjustment for cell composition (covariate set **A**) there were 169,951 loci significantly associated with chronological age. The original publication reported 137,993 significant loci [15].

## Discussion

We trained and evaluated a model that estimates the proportions of fifteen leukocyte subtypes in a large cohort that had both whole blood DNAm microarray and CBCs/flow cytometry data. We demonstrated in a large test set that the estimates generated from our model explain more of the variance observed in whole blood DNAm levels and can vastly reduce the number of false-positive findings in epigenome-wide association studies. In many situations, researchers may benefit by replacing existing reference-based cell deconvolution models that provide estimates for six to seven cell types with our extended model.

One advantage of our model is that by training on a large subset of HRS it should be valid for diverse study populations over the age of 50. In comparison, many existing cell-type deconvolution models follow the Houseman method of using reference datasets of purified cell types [4]. These reference datasets are typically limited in the number and demographics of the sample donors (Reinius et al. [3]: 6 healthy Swedish male donors aged 25-60; Salas et al. [16]: 31 males and 6 females aged 19–59, all healthy). The subset of HRS used here included 2530 individuals who were 50-100 years old, racially diverse, and of good and bad health. A major strength of our study is this 100-fold increase in the number of sample donors, resulting in better representation. Future studies may demonstrate generalizability to younger individuals and ultimately even within the given age range, as our training and test datasets are from the same underlying HRS population. We applied our model to an external dataset featuring a broad age range (14-94 years) and showed that our model substantially lowered genomic inflation and generated far fewer age-related EWAS hits, demonstrating the utility of our cell proportion estimates. A comparison with a newer 12-cell type model (requiring a licensing agreement) is planned for a future version of this manuscript [17].

An obvious benchmark of model performance would be to assess the correlation between estimated and measured cell proportions (we report these in Table 2). However, this approach is flawed without a gold standard or if said standard comes with substantial measurement error. As with all laboratory measures, there may be measurement error in flow cytometry counts [18]. Instead, we assess model performance by quantifying how much of the variance in whole blood DNAm levels is explained by the various cell proportion estimates. In a similar project, we built a model to infer proportions of five available cell types, i.e., neutrophils, eosinophils, basophils, monocytes, and lymphocytes [19]: correlation coefficients between estimated and measured proportions in the test set ranged from 0.02 (basophils) to 0.85 (neutrophils). Yet estimated cell proportions in a test set explained more of the variance in methylation levels than measured proportions on a per-loci basis. We observe the same pattern here (see Figure 2). There are two possible explanations. In the first scenario, cell proportion estimates, because they are derived from a subset of the DNAm data, might (unintentionally) capture variance beyond that associated with cell composition, such as batch effects or latent factors. Plugging those estimates back into regression models could, therefore, explain a larger share of variance in methylation levels even if the estimates themselves were less precise. Alternatively, the estimates could be more precise than the measured cell proportions. In this scenario, the correlation between estimates and measured proportions would be attenuated due to measurement error. The model could produce estimates that exceed the precision of the input variables as the model parameters are averages across many samples and predictions are made based on redundant features. Naturally, a combination of both scenarios is possible. However, we took further precautions to reduce the impact of batch effects by calculating the differences in R^2^ from a baseline model that included sex, age, plate, and principal components of quality control metrics.

As a whole, the estimated cell composition from our new model explained 14.1p of the variance compared to 9.5pp when including the same number of covariates from the measured cell composition, a difference of 4.5pp. This statistic was calculated on the HRS test set and excluded any loci used as features in our model. Our model also performs better than estimates based on the Reinius reference panel [3] which explained 10.7pp in the same test set. Exact values will vary between study populations and flow cytometry instruments, and researchers may use the proportion of variance explained to guide their decision of which set of cell-type proportions will be the best choice for their analyses.

In case a benchmark design as discussed above is not possible, we prefer correlation coefficients between estimated and measured cell proportions as opposed to root mean squared errors (which we include in Table 2) as the former are the better indicators of how well these estimates can adjust for confounding when added as covariates to linear regression models.

We picked the top 50 markers for each cell type during model training. As the predictive power of these markers should be redundant, our model should perform well even if some features are missing for a particular sample. More sophisticated ways of selecting features may improve model performance, something we did not investigate.

In this study, we developed a deconvolution algorithm for estimating proportions of fifteen leukocyte subtypes from DNAm data and provide an implementation in an R package. Cell composition is a major contributor to variance in DNA methylation measures, and cell composition can be operationalized in epidemiologic frameworks in multiple ways including as a precision variable and as a mediator [20]. The success of EWAS depends on adjustment, most notably for batch effects, and cell composition, and our model facilitates the latter.

## Abbreviations

CBC: complete blood count
CPT: cryopreservation tube
DNAm: DNA methylation
EWAS: epigenome-wide association study
HRS: Health and Retirement Study
pp: percentage points

## Data and code availability statement

HRS epigenetic data can be accessed via the National Institute on Aging Genetics of Alzheimer’s Disease Data Storage Site (NIAGADS). HRS flow cytometry data are also publicly available from the HRS website at https://hrs.isr.umich.edu/. The code to produce the analyses in this paper is available on GitHub and Zenodo (github.com/hhhh5/HRS/, doi.org/10.5281/zenodo.7369326). Cell type deconvolution for Illumina 450k and EPIC microarrays using the model trained here is implemented in the open-source *ewastools* R package (github.com/hhhh5/ewastools, doi.org/10.5281/zenodo.7329019) by the *estimateLC* function and can be used without restrictions (public domain).

## Acknowledgments

These analyses were supported by NIH grants P30 ES023515, R00 ES023450, and R01 AG067592. The Health and Retirement Study is sponsored by the National Institute on Aging (grant U01AG009740) and is conducted by the University of Michigan. We thank the participants and staff of the Health and Retirement Study.

